# Innervation modulates the functional connectivity between pancreatic endocrine cells

**DOI:** 10.1101/2020.11.04.368084

**Authors:** Yu Hsuan Carol Yang, Linford J.B. Briant, Christopher Raab, Sri Teja Mullapudi, Hans-Martin Maischein, Koichi Kawakami, Didier Y.R. Stainier

**Affiliations:** Department of Developmental Genetics, Max Planck Institute for Heart and Lung Research, Bad Nauheim, Germany; Oxford Centre for Diabetes, Endocrinology and Metabolism, Radcliffe Department of Medicine, University of Oxford, Oxford, United Kingdom; Division of Molecular and Developmental Biology, National Institute of Genetics, Department of Genetics, SOKENDAI, Mishima, Japan

## Abstract

Direct modulation of pancreatic endocrine cell activity by autonomic innervation has been debated. To resolve this question, we established an *in vivo* imaging model which also allows chronic and acute neuromodulation. Starting at a stage when zebrafish islet architecture is reminiscent of that in adult rodents, we imaged calcium dynamics simultaneously in multiple islet cell types. We first find that activity coupling between beta cells increases upon glucose exposure. Surprisingly, glucose exposure also increases alpha-alpha, alpha-beta and beta-delta coordination. We further show that both chronic and acute loss of nerve activity diminish activity coupling, as observed upon gap junction depletion. Notably, chronic loss of innervation severely disrupts delta cell activity, suggesting that delta cells receive innervation which coordinates its output. Overall, these data show that innervation plays a vital role in the establishment and maintenance of homotypic and heterotypic cellular connectivity in pancreatic islets, a process critical for islet function.

## Introduction

Tight regulation of hormone release from pancreatic islets is critical for glucose homeostasis and its disruption can lead to diabetes mellitus^1^. Pancreatic islets are composed of different cell types, including the hormone producing alpha, beta and delta cells, peripheral nerves, and vascular endothelial and smooth muscle cells. Studies have implicated signaling from the vascular scaffold^2,3^ and nerve networks^4–9^ during the development and function of pancreatic islet cells. However, it remains difficult to investigate the immediate effects of acute nerve modulation on islet cell function. Given the alterations in islet innervation architecture in some models of diabetes^10–12^, it is imperative to understand whether disruption of nervous control can contribute to diabetes etiology.

Different methods of assessing islet cell function have provided important clues into the role of autocrine and paracrine signaling in this process. Electrophysiological recordings have provided fundamental insights into isolated islet cell function^13–15^, including functional connectivity studies that identified homotypic as well as heterotypic coupling between endocrine cells^16,17^. However, assessing islet function in live animals with undisrupted vascular and nerve networks remains challenging. Calcium dynamics is a good readout of the function of all islet cell types because its influx is critical for hormone release. However, no studies to date have been able to record simultaneously the activity of all islet endocrine cell types in the intact organ of a living animal, which is required to understand how the different endocrine cell types respond to physiological perturbations individually and interdependently. To this end, we established an *in vivo* imaging platform to visualize the activity of all the islet cell types by combining calcium imaging with cell type reporters. We investigated the functional connectivity between homotypic and heterotypic cell pairs by analyzing the correlation patterns in their intracellular calcium changes. Chronic and acute inhibition of nerve activity captured its dynamic control of the functional connectivity between islet endocrine cells, which is in part dependent on gap junctions.

## Results and discussion

### The activity of all pancreatic endocrine cell types can be studied simultaneously in vivo

The zebrafish primary islet becomes highly innervated^7^ and vascularized^3,18,19^ early in development (Figure 1A). Fluorescent reporters for different pancreatic endocrine cell types, including beta, alpha, and delta cells, were used to study the establishment of islet cytoarchitecture (Figure 1B). By 100 hours post fertilization (hpf), a beta cell core and alpha cell mantle layout is observed (Figure 1B-C), reminiscent of adult rodent islets^20^ and small human islets^21^. Simultaneous functional assessment of all islet cell types *in vivo* required reporters for cell activity and cell identity. We used the *Et(1121A:GAL4FF)* enhancer trap line with the *Tg(UAS:GCaMP6s)* line for calcium imaging of all islet cells (Figure 1D-E, Figure 1-Movie Suppl. 1), as well as the *Tg(ins:mCardinal)* and *Tg(sst2:RFP)* lines to assign beta and delta cell identity, respectively; alpha cells were identified by their mantle localization and/or by immunostaining (Figure 1E). Thus, for the first-time, we were able to assess the activity of all islet cells in their native environment in an intact living animal (Figure. 1F, Figure 1 - Movie Suppl. 2).

**Figure 1.**
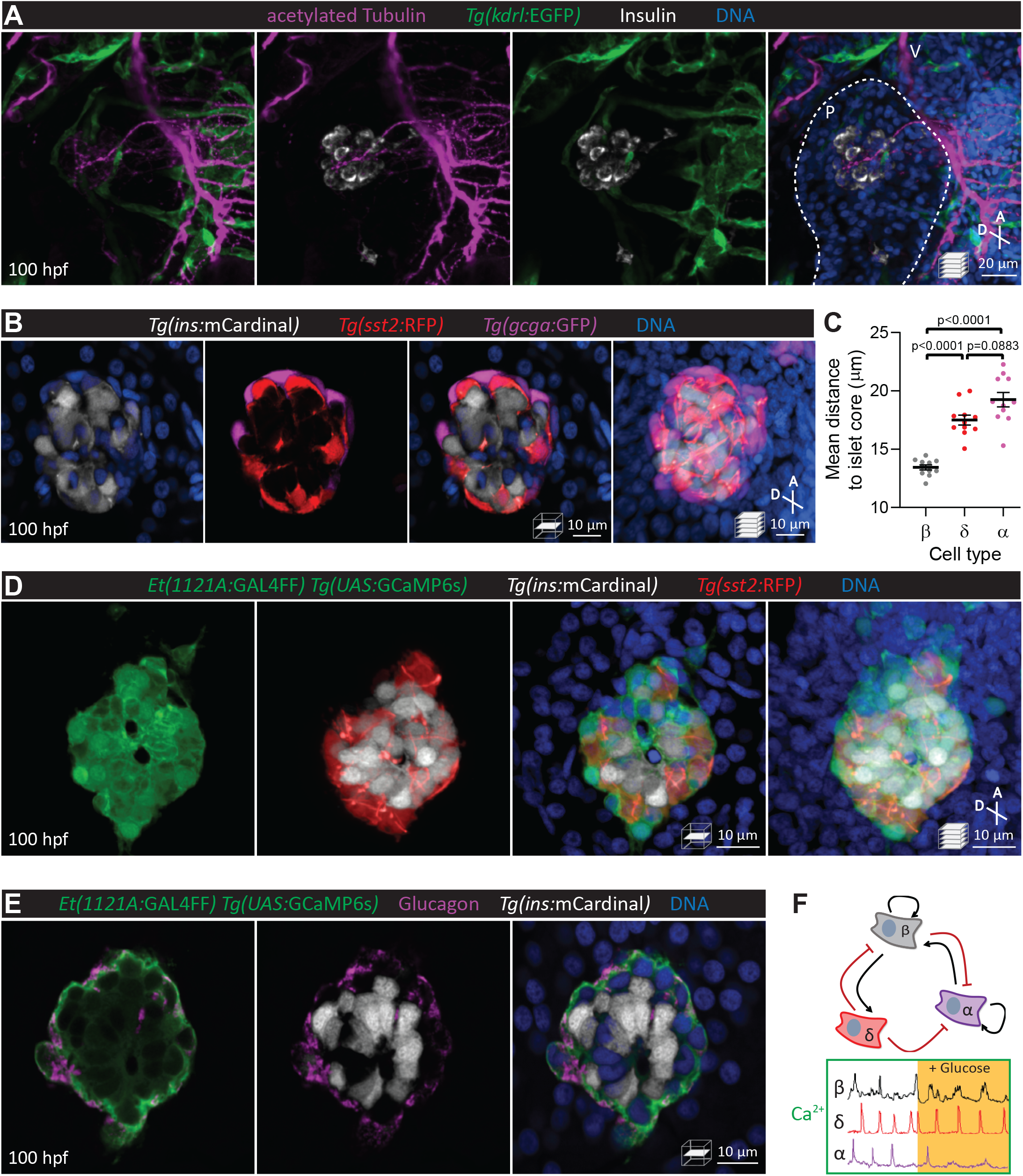
Pancreatic islet cell activity is visualized *in vivo* with preserved vascular and neural networks. **A.** Wholemount immunostaining of wild-type zebrafish at 100 hours post fertilization (hpf) for acetylated Tubulin (nerves), *Tg(kdrl:*GFP) expression (vessels), and Insulin (beta cells), and counterstaining with DAPI (DNA). **B.** 100 hpf *Tg(ins:mCardinal); Tg(sst2:RFP); Tg(gcga:GFP)* zebrafish stained with DAPI (DNA). **C.** Mean distance of pancreatic islet cells to islet core reveals a beta cell core and alpha cell mantle; mean ± S.E.M., n=11 animals; p-values from ANOVA. **D.** 100 hpf *Et(1121A:GAL4FF); Tg(UAS:GCaMP6s); Tg(ins:mCardinal); Tg(sst2:RFP)* zebrafish stained with DAPI (DNA). **E.** 100 hpf *Et(1121A:GAL4FF); Tg(UAS:GCaMP6s); Tg(ins:mCardinal)* zebrafish stained for Glucagon (alpha cells) and DNA. **F.** Schematic of documented interactions between beta, delta, and alpha cells and of intracellular calcium recordings in each of these cell types. Maximum intensity projections or single planes are presented; A, anterior; D, dorsal; V, vagus nerve; P, pancreas.

### Homotypic and heterotypic coupling between endocrine cells requires pancreatic innervation

Activity coupling between pancreatic endocrine cells can be mediated by autocrine and paracrine signaling, gap junctions, and other means. To investigate whether pancreatic innervation is critical for intra-islet coordination of activity, we used different approaches to chronically or acutely inhibit neural signaling. We used endoderm transplantation to generate chimeric zebrafish that express two GAL4/UAS systems in different germ layer-derived tissues and investigated the role of chronic neural inhibition on islet function (Figure 2A). Pan-neural expression of botulinum toxin (BoTx) chronically inhibits neurotransmitter release^7,22^ and leads to elevated glucose levels at 100 hpf^7^ (Figure 2B). While primary islet volume was consistently greater in BoTx^+^ larvae at 100 hpf (Figure 2C; as we have described for earlier stages^7^), we did not observe changes in the architectural arrangement of the different islet cell types (Figure 2D). From the normalized single cell calcium traces, the correlation matrices, and average correlation coefficients (Ravg), we observed that the calcium dynamics in BoTx^+^ larvae were significantly disrupted, with impairment in beta cell coupling (Figure 2E-G). Our measure of Ravg over increasing intercellular distance also revealed severe disruption in BoTx^+^ larvae, indicating drastically altered synchronicity and wave propagation (Figure 2H-I). To determine how dysfunctional neural signaling influenced communication between different endocrine cell types, we conducted fraction time analysis of heterotypic cell pairs that were in nearest proximity to each other. We reasoned that nearest neighbours have a greater likelihood of displaying coupling, which would be evident by analyzing activity patterns between heterotypic cell pairs. We analyzed the single cell calcium traces and determined the fraction of time a given cell pair resides in a state when 1) one cell was silent, the other active, 2) both cells were active, and 3) both cells were silent. This analysis provided insights into the neural regulation of heterotypic intercellular communication. Aside from the quiet phase, when both cell types were silent, we found changes in activity patterns between delta-beta, alpha-delta, and alpha-beta cell pairs (Figure 2J-L): upon chronic neural inhibition, the delta-silent/beta-active state was decreased while the delta-active/beta-silent state (Figure 2J) and the delta-active/alpha-silent state (Figure 2K) were both increased. Overall, chronic neural inhibition resulted in the hyperactivation of delta cells, and somatostatin released from delta cells likely inhibited both beta and alpha cell activity. We also found a significant decrease in the alpha-active/beta-active state upon chronic neural inhibition (Figure 2L). With growing evidence of local glucagon signaling in potentiating beta cell activity^23^, this finding further suggests that neural regulation may be an important regulator of alpha-beta connectivity.

**Figure 2.**
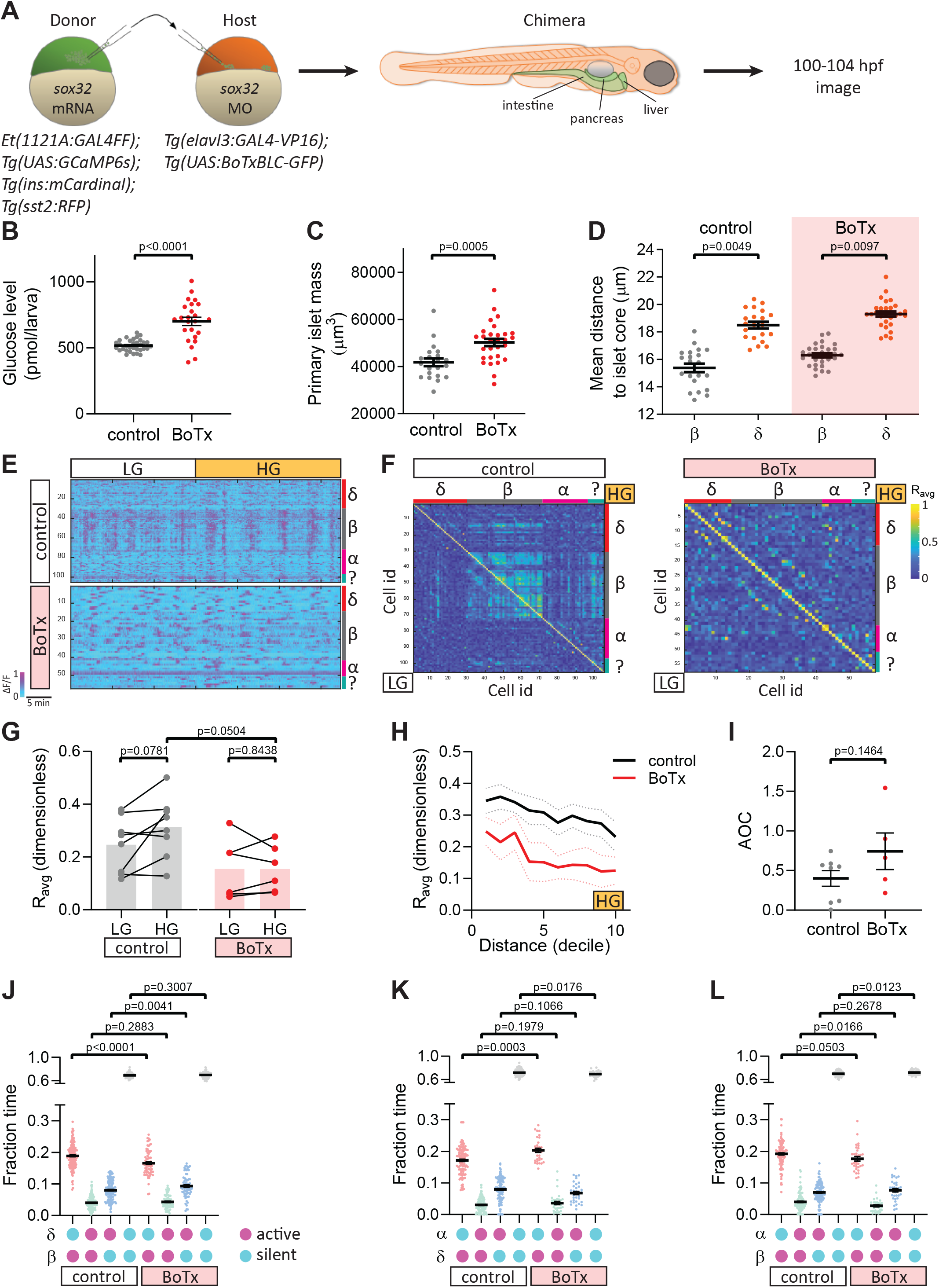
Chronic inhibition of synaptic transmission disrupts islet cell activity. **A.** Schematic of transplants to generate chimeras with endodermal organs derived entirely from donor embryos. **B.** Whole larva free glucose level measurements at 100 hpf, mean ± S.E.M., n=24-32 batches of 5 larvae per replicate. **C.** Quantification of primary islet mass. **D.** Mean distance of pancreatic islet cells to islet core, mean ± S.E.M., n=21-29 animals, p-values from t-tests. **E.** Normalized calcium traces of pancreatic islet cells (including delta, beta, alpha, and unidentified cells). Individual islet cells were assigned to a cell type and given a cell id; LG, basal condition; HG, glucose treated condition. **F.** Correlation matrices of islet cell activity. Individual islet cells were assigned to a cell type and given a cell id, and average correlation coefficients for given cell pairs (matrix row-column intersects) were calculated for LG (basal condition; bottom left triangle) and HG (glucose treated condition; top right triangle). **G.** Average beta cell correlation coefficients in individual larvae, n=5-8 animals, p-values from t-tests. **H.** Average beta cell correlation coefficients with cell distance distribution from 1 (close) to 10 (far), mean ± S.E.M. (dotted lines). **I.** Area over the curve (AOC) analysis of H, mean ± S.E.M., n=5-8 animals, p-values from t-tests. **J-L.** Fraction time analysis of heterotypic delta-beta **(J)**, alpha-delta **(K)** and alpha-beta **(L)** cell pairs, mean ± S.E.M., n=32-160 cell pairs; magenta circle, active state; cyan circle, silent state.

Given the potential role of pancreatic innervation on islet cell maturation, we next investigated the effects of acutely blocking neural activity using two different approaches. By lineage tracing, we found that the neural crest derived peri-islet neurons were also labelled by the *Et(1121A:GAL4FF)* enhancer trap (Figure 3A-B), thereby allowing us to investigate the effects of photo-ablating a subset of peri-islet neurons on islet cell activity (Figure 3C). While the oscillatory pattern of calcium dynamics was maintained in beta cells upon this ablation (Figure 3D), the coupling between beta cells was significantly decreased (Figure 3E-F). Notably, we did not observe further impairment in beta cell wave propagation (Figure 3G-H), suggesting that upon ablation of peri-islet neurons, the signal that initiated beta cell waves was blunted while the beta cells maintained their propensity for calcium wave propagation. Unlike in the chronic neural inhibition scenario, delta-beta and alpha-delta coupling was not affected upon ablating peri-islet neurons (Figure 3I-J). However, we observed a trend towards a decrease in the alpha-active/beta-active state (Figure 3K). Overall, ablation of peri-islet neurons significantly disrupted beta-beta connectivity, while conclusions regarding heterotypic interactions in this model will require further investigations into the various neural subsets that were targeted. It is likely that our targeting of peri-islet neurons did not always ablate the specific neurons that guide the activity coupling between alpha and beta cells.

**Figure 3.**
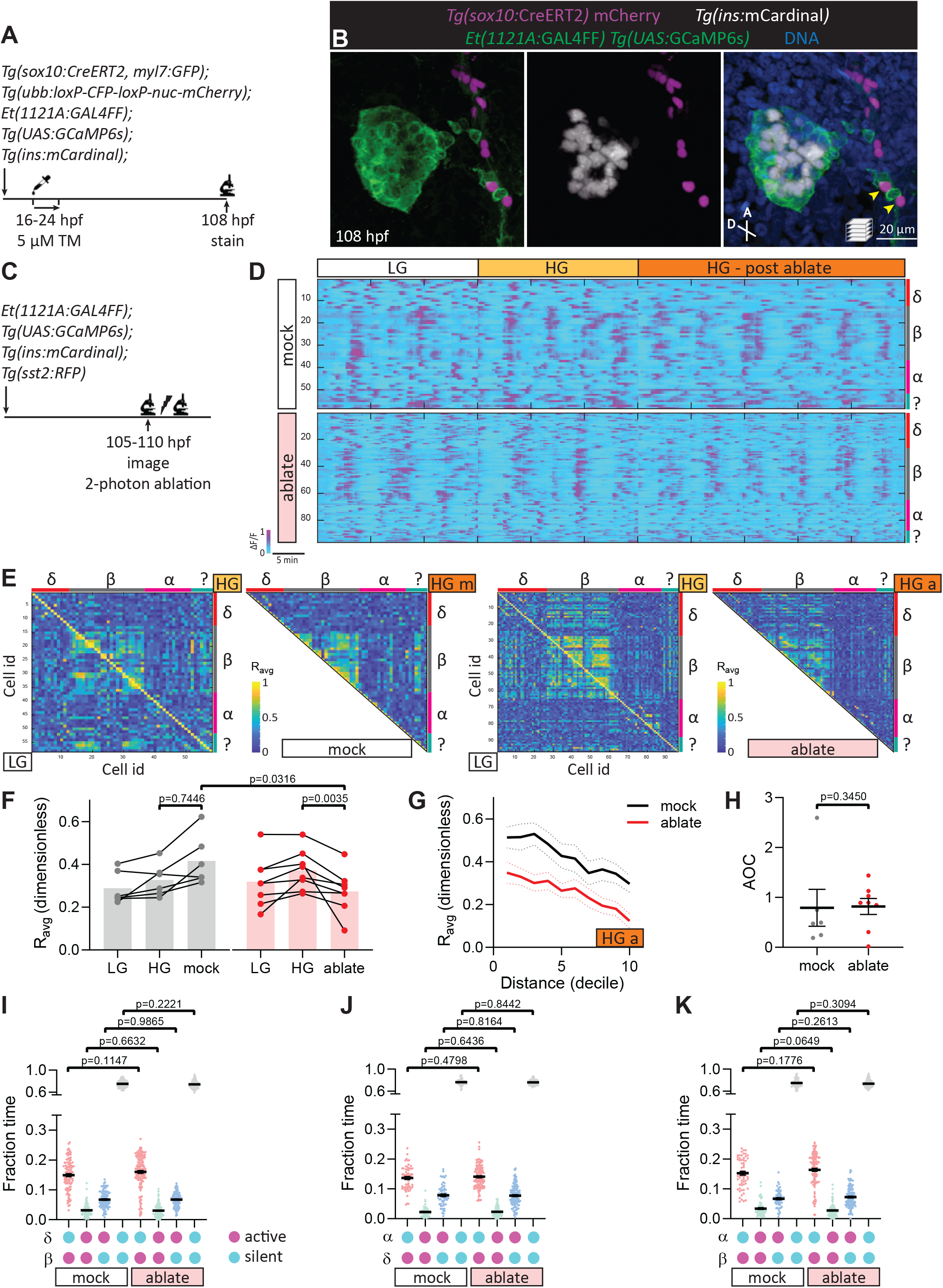
Targeted ablation studies reveal the crucial role of peri-islet neurons for islet cell activity. **A.** Schematic of lineage tracing of neural crest cells in *Tg(sox10:CreERT2, myl7:GFP); Tg(ubb:loxP-CFP-loxP-nuc-mCherry); Tg(ins:mCardinal); Et(1121A:GAL4FF); Tg(UAS:GCaMP6s)* zebrafish following 5 μM tamoxifen (TM) treatment from 16 to 24 hpf and staining at 108 hpf. **B.** Wholemount immunostaining at 100 hpf for mCherry expression (neural crest derived cells) and counterstaining with DAPI (DNA). Yellow arrowheads point to neural crest cells positive for GCaMP6s expression. **C.** Schematic of two-photon ablation experiment. **D.** Normalized calcium traces of pancreatic islet cells (including delta, beta, alpha, and unidentified cells). Individual islet cells were assigned to a cell type and given a cell id; LG, basal condition; HG, glucose treated condition. **E.** Correlation matrices of islet cell activity; LG, basal condition; HG, glucose treated condition. **F.** Average beta cell correlation coefficients in individual larvae, n=6-8 animals, p-values from t-tests. **G.** Average beta cell correlation coefficients with variable cell distance, mean ± S.E.M. (dotted lines). **H.** Area over the curve (AOC) analysis of G, mean ± S.E.M., n=5-8 animals, p-values from t-tests. **I-K.** Fraction time analysis of heterotypic delta-beta **(I)**, alpha-delta **(J)** and alpha-beta **(K)** cell pairs, mean ± S.E.M., n=63-145 cell pairs; magenta circle, active state; cyan circle, silent state.

Next, we took an optogenetic approach by generating a transgenic line that allows one to acutely photo-inhibit the release of neurotransmitters with a single pulse of blue light. This method has previously been used in *Drosophila*^24^ and *C. elegans*^25,26^ for targeted photo-ablation and photo-inhibition. Pan-neural expression of a singlet oxygen generator, miniSOG2, tethered to synaptic granules resulted in a blue light inducible loss of swimming activity in 110 hpf larvae (Figure 4 – Fig. Suppl. 1A-C). Following this confirmation of the effectiveness of the tool, we studied neural control of islet cell activity upon acute photo-inhibition (Figure 4 – Fig. Suppl 1D). Surprisingly, photo-inhibition decreased glucose levels compared to transgene-negative fish exposed to the same light condition (Figure 4A). Similar to peri-islet neural ablation, pan-neural photo-inhibition decreased beta cell connectivity (Figure 4B-D), while beta cell wave propagation was not impaired (Figure 4E-F). Changes in delta-beta and alpha-beta heterotypic interactions was observed upon acute neural inhibition (Figure 4G-I). A significant decrease in delta-silent/beta-active state reflects what we observed upon chronic neural inhibition (Figure 4G). Like photo-ablation of peri-islet neurons, no changes in alpha-delta interactions were observed (Figure 4H). Notably, following acute neural inhibition both alpha-silent/beta-active and alpha-active/beta-active states were significantly decreased, while alpha-active/beta-silent and alpha-silent/beta-silent states were significantly increased (Figure 4I). These changes in alpha-beta interactions were consistently observed upon both chronic and acute pan-neural inhibition, possibly reflecting a role for neurons in the maintenance of alpha-beta coupling.

**Figure 4.**
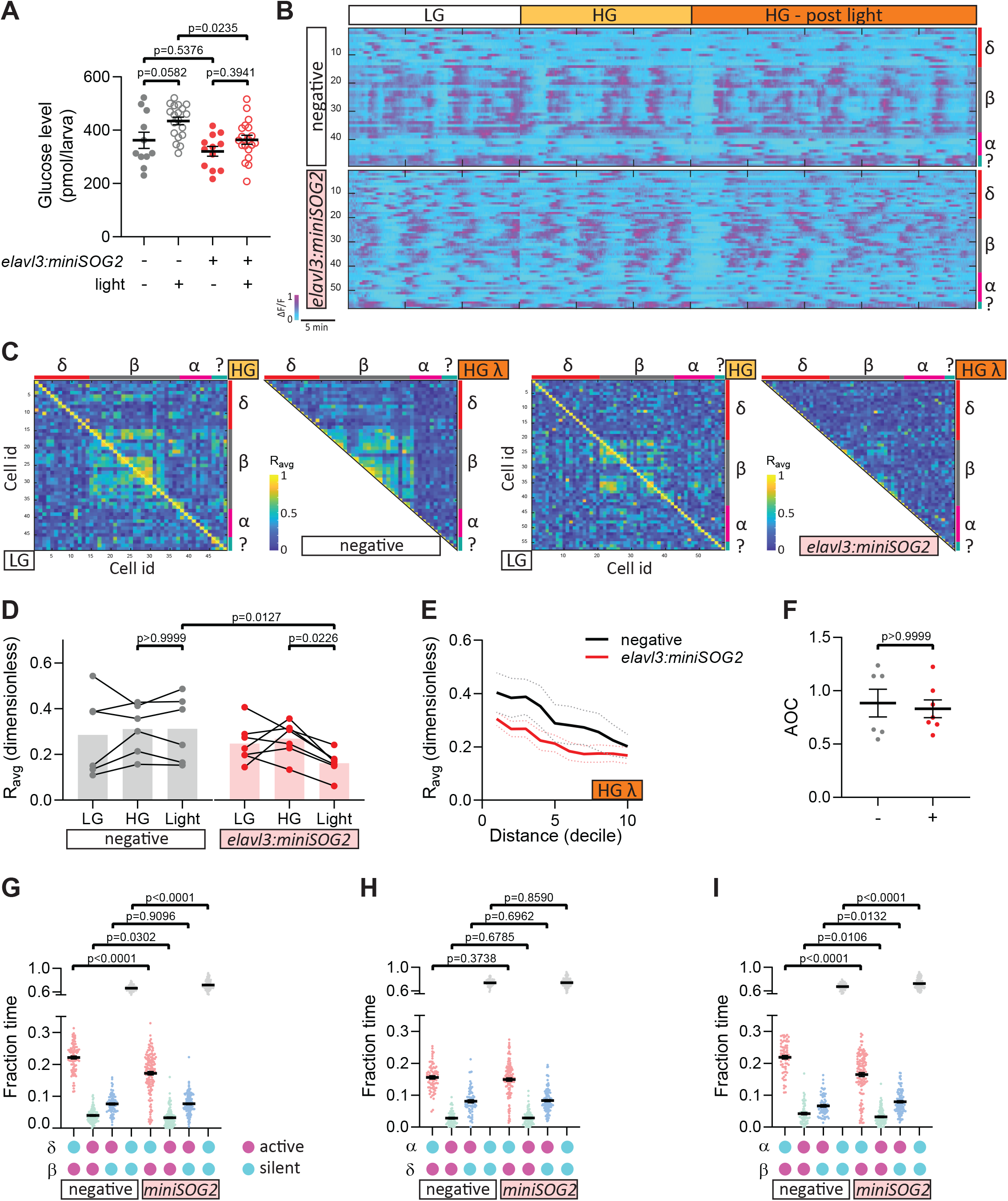
Acute optogenetic inhibition of neurotransmitter release disrupts islet cell activity. **A.** Whole larva free glucose level measurements of *Tg(elavl3:sypb-miniSOG2-P2A-mScarlet)* zebrafish at 110 hpf, mean ± S.E.M., n=11-19 batches of 5 larvae per replicate. **B.** Normalized calcium traces of pancreatic islet cells (including delta, beta, alpha, and unidentified cells). Individual islet cells were assigned to a cell type and given a cell id; LG, basal condition; HG, glucose treated condition. **C.** Correlation matrices of islet cell activity; LG, basal condition; HG, glucose treated condition. **D.** Average beta cell correlation coefficients in individual larvae, n=6-7 animals, p-values from t-tests. **E.** Average beta cell correlation coefficients with variable cell distance, mean ± S.E.M. (dotted lines). **F.** Area over the curve (AOC) analysis of E, mean ± S.E.M., n=6-7 animals, p-values from t-tests. **G-I.** Fraction time analysis of heterotypic delta-beta **(G)**, alpha-delta **(H)** and alpha-beta **(I)** cell pairs, mean ± S.E.M., n=71-119 cell pairs; magenta circle, active state; cyan circle, silent state.

### Gap junctions are required for beta cell coupling

Could innervation act as the upstream trigger for activity coupling while other means of intercellular communication propagate the signals across the islet? Gap junctions have well described roles in mediating coordination of beta cell activity and insulin secretion^27,28^. In mammals, *cx36* encoded gap junctions mediate the majority of the functional coupling between beta cells^29–31^. While previous reports have described the lack of beta cell expression of the zebrafish homolog *cx35*^32^, we found that *cx43* is highly expressed in zebrafish beta cells by 100 hpf (Figure 5A, Figure 5 – Fig. Suppl. 1). Connexin 43 has known roles in mediating bone growth and regeneration in zebrafish^33–35^, however no pancreatic defects have been reported in these loss-of-function studies. We used animals harboring a *cx43* mutation in the exon encoding the second extracellular loop and that has been proposed to interfere with connexon docking between neighbouring cells^36^, to elucidate the role of *cx43* in beta cells (Figure 5B). While *cx43^+/−^* and *cx43^−;/−^* larvae do not exhibit gross phenotypic defects at 100 hpf (Figure 5 – Fig.Suppl. 2A-B), a subset of them (23.5% and 17.9%, respectively) display elevated glucose levels (Figure 5 – Fig. Suppl. 2C), which is likely an underestimation of the true penetrance of this phenotype due to the pooling of animals required for the assay. Since islet architecture in *cx43^+/−^* and *cx43^−;/−^* larvae at 100 hpf appeared unaffected (Figure 5C), we next assessed the activity coupling between beta cells. From normalized single cell calcium traces, we observed blunted glucose-induced coupling of beta cell activity in *cx43^+/−^* and *cx43^−;/−^* larvae (Figure 5D, Figure 5 – Movie Suppl. 1). This finding was confirmed in the correlation matrices (Figure 5E) and average correlation coefficients (R_avg_; Figure 5F). While we did not observe a complete penetrance of this phenotype, there was a significant decrease in R_avg_ following high glucose treatment in *cx43^+/−^* larvae (Figure 5F). Given the reduced coupling between beta cells in the entire islet, we next analyzed whether coupling changes relative to intercellular distance. The greater decrease in R_avg_ with increasing cellular distance observed in *cx43^−;/−^* larvae (Figure 5G-H) indicates that the propagation of calcium waves across the islet was significantly impaired upon loss of Cx43 function. Given that some coupling was maintained between beta cells in *cx43^−;/−^* larvae, it is likely that pancreatic innervation could be mediating the remaining intra-islet coordination of activity.

**Figure 5.**
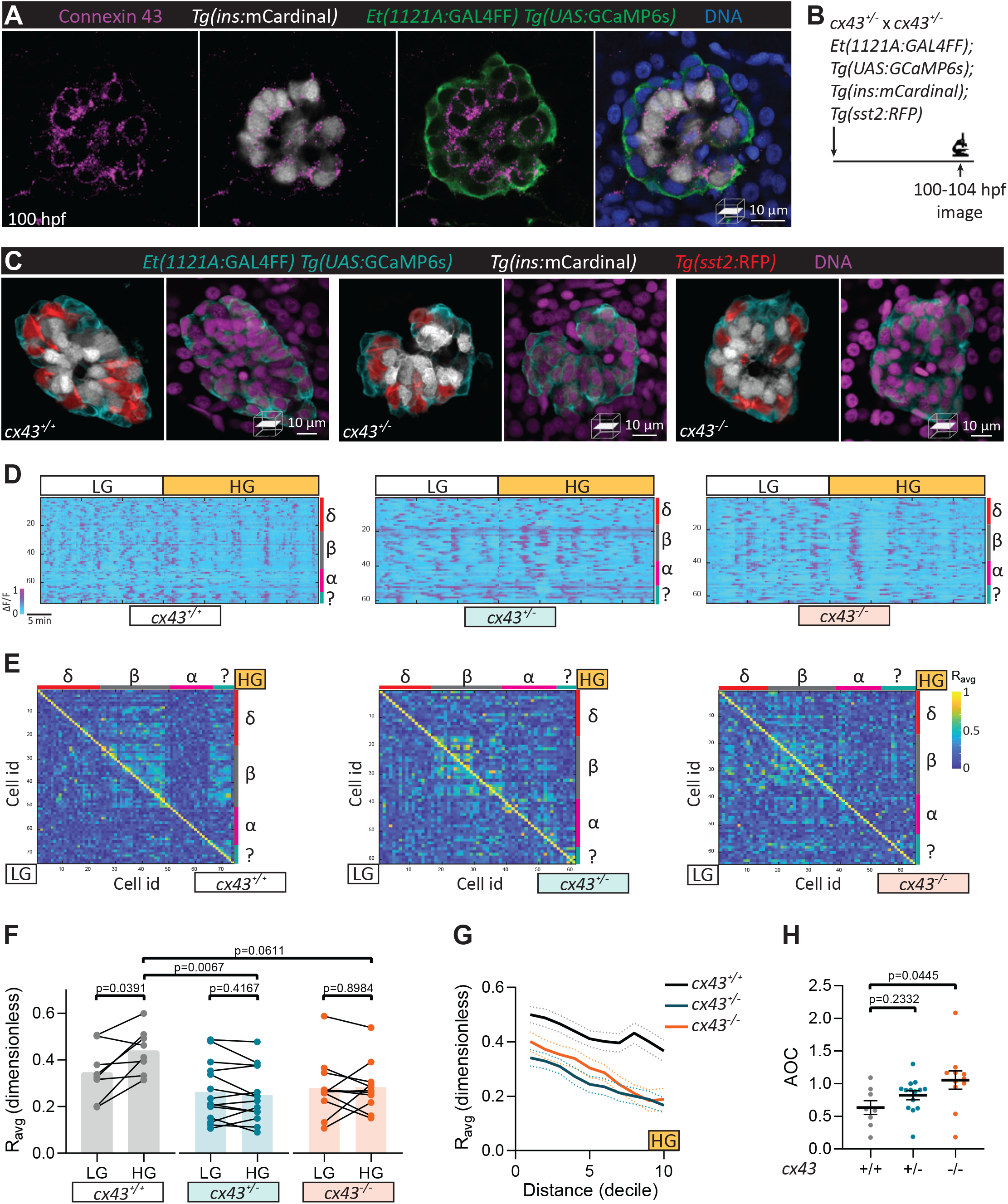
Coordination of beta cell activity is in part mediated by Cx43 gap junctions. **A.** Wholemount immunostaining of 100 hpf *Et(1121A:GAL4FF); Tg(UAS:GCaMP6s); Tg(ins:mCardinal)* zebrafish for Connexin 43, and counterstaining with DAPI (DNA). **B.** Schematic of *cx43* mutant experiment. **C.** *Et(1121A:GAL4FF); Tg(UAS:GCaMP6s); Tg(ins:mCardinal); Tg(sst2:RFP) cx43^+/+^*, *cx43^+/−^*, and *cx43^−;/−^* larvae imaged at 100 hpf. **D.** Normalized calcium traces of pancreatic islet cells (including delta, beta, alpha, and unidentified cells). Individual islet cells were assigned to a cell type and given a cell id; LG, basal condition; HG, glucose treated condition. **E.** Correlation matrices of islet cell activity. **F.** Average beta cell correlation coefficients in individual larvae, n=8-14 animals, p-values from t-tests. **G.** Average beta cell correlation coefficients with variable cell distance, mean ± S.E.M. (dotted lines), n=8-14 animals. **H.** Area over the curve (AOC) analysis of G, mean ± S.E.M., n=8-14 animals, p-values from ANOVA.

Fraction time analysis of heterotypic cell pairs determined how dysfunctional Cx43 gap junctions influenced the other islet cell types (Figure 5 – Fig. Suppl. 2D-F). While no significant changes were observed between alpha-delta and alpha-beta activity patterns, a significant increase in delta-silent/beta-active state was observed in *cx43^+/−^* and *cx43^−;/−^* larvae (Figure 5 – Fig. Suppl. 2D). Given that *cx43* expression is weakly detected in delta cells, it may also be important in mediating the communication between delta and beta cells. Indeed, gap junction mediated beta to delta electrical coupling has been proposed to activate glucose induced delta cell activation, at least in part^17^, and we found a significant decrease in the delta-active/beta-active state in *cx43^+/−^* larvae (Figure 5 – Fig. Suppl. 2D). Interestingly *cx43* expression is absent in adult beta and delta cells^37^ (Figure 5 – Fig. Suppl. 1), yet we observed severe fasting hypoglycemia in *cx43^+/−^* and *cx43^−;/−^* adults (Figure 5 – Fig. Suppl. 2G). Whether dysfunctional Cx43*-*mediated gap junctional coupling between beta-beta and delta-beta cells early in life influences glucose homeostasis in adults warrants further studies.

Dissecting the complex interplay of local autocrine, paracrine, and gap junctional communication between different endocrine cells, in addition to vascular and nerve interactions, is often hindered by our ability to simultaneously study them in an intact organ within its innate environment. Imaging calcium dynamics with genetically encoded biosensors or calcium sensitive fluorescent indicators in individual islet cell types has been conducted *in vitro* with dispersed cells^38,39^, whole islets^40,41^, and perfused pancreas slices^42,43^, and *in vivo* with islets transplanted into the anterior chamber of the eye^4,44^, and intravital imaging of mouse pancreas^45^. We report a non-invasive *in vivo* imaging strategy to study all the different pancreatic endocrine cell types within the same animal. Our three approaches to inhibit neural control, ranging in temporal and spatial specificity, provided useful insights into the role of neurons in regulating pancreatic islet function. Activity coupling between beta cells is in part mediated by Cx43 gap junctions, and neural regulation is critical for the establishment and maintenance of beta cell connectivity as we consistently found decreased beta cell coupling upon chronic and acute neural inhibition. Given that our targeted neural ablation approach also led to this decline in beta cell coupling, it is likely that autonomic neural control is required for beta cell connectivity independently of indirect effects resulting from neural regulation of other organs. It has been proposed that glucose sensing neurons regulate early postnatal beta cell proliferation and maintenance of beta cell function^9^, and our data support the peri-islet localization of such neurons.

We have focused on the pancreatic beta, alpha, and delta cells; however, it is important to note that there are other endocrine cell types (gamma and epsilon cells) that remain undefined in our study, but do display glucose induced activity coupling with beta cells (Figure 2F). Fraction time analysis allowed us to study activity patterns of heterotypic cell pairs that are near each other upon loss of Cx43 or neural signaling. We found changes in delta-beta activity patterns, supporting a role for gap junctions in mediating electrical coupling between delta and beta cells^17^. Both chronic and acute neural inhibition blunted the functional connectivity between alpha and beta cells suggesting that neurons may have an important role in the paracrine potentiation of beta cell activity by glucagon released from alpha cells^23^. We propose that alpha-beta interactions are specifically regulated by pancreatic innervation, since peri-islet neural ablation also led to a similar decreasing trend in the alpha-active/beta-active state. However, given that pan-neural inhibition is required to induce changes in delta-beta interactions, it is possible that central nervous control of other organs could in part be driving these changes. Importantly, our data further support the role of neurons in modulating delta cell activity and maturation, given that hyperactivity likely reflects uncontrolled release of somatostatin and the delta cell hypertrophy and decreased delta cell mass we previously described^7^.

Selective targeting of subsets of neurons will advance our understanding of the pancreatic islet-neural interplay in health and disease, including diabetes pathophysiology. Through our *in vivo* studies of homotypic and heterotypic activity coupling, we have illustrated how studying functional connectivity can be achieved for the endocrine pancreas and discovered a critical role for neurons in mediating these connections. Given the cellular heterogeneity of organ composition, simultaneously evaluating the function of the different cell types that make up an organ provides insights that can be missed by simply investigating one cell type at a time. Interrogating the neural regulation of other organs, such as the liver, intestine, and kidney, can be achieved through the extended application of these tools. In combination with the ability to monitor organ development, function, and regeneration *in vivo,* this approach will allow one to address difficult questions pertaining to the autonomic nervous system and its role in organ maintenance and dysfunction.

## Materials and methods

### Zebrafish transgenic lines and husbandry

All zebrafish husbandry was performed under standard conditions in accordance with institutional (MPG) and national ethical and animal welfare guidelines. Adult zebrafish were fed a combination of fry food (Special Diet Services) and brine shrimp five times daily and maintained under a light cycle of 14 hours light: 10 hours dark at 28.5°C. Zebrafish embryos and larvae were grown in egg water at 28.5°C. Transgenic and mutant lines used were on the *mitfa^w2/w2^* background and as described in Table 1.

**Table 1.**
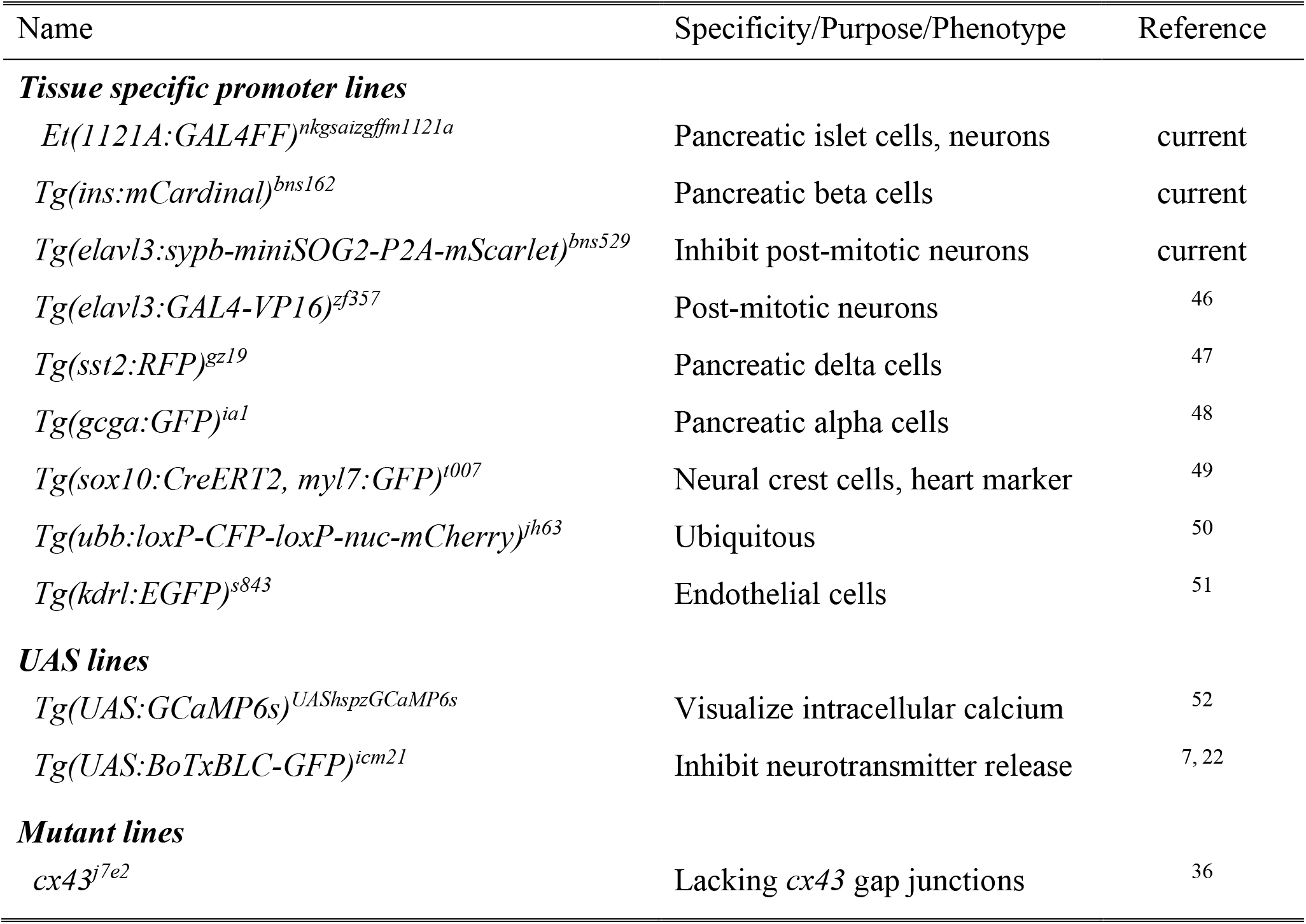
List of zebrafish transgenic and mutant lines.

*Tg(elavl3:sypb-miniSOG2-P2A-mScarlet)* line was generated by Tol2 transgenesis of 8.7 kb *elavl3* promoter (Addgene: 59531, AgeI restriction enzyme digested) driving expression of *sypb* (Addgene: 74316), *miniSOG2* (Addgene: 87410), *P2A-mScarlet* (gift from A. Beisaw) cloned with Cold Fusion (System Biosciences). *Tg(ins:mCardinal)* line was generated by Tol2 transgenesis of 1.1 kb *ins* promoter (in-house plasmid, MluI restriction enzyme digested) driving expression of *mCardinal* (Addgene: 51311) cloned with Cold Fusion.

### Generation of chimeric zebrafish

*Et(1121A:GAL4FF); Tg(UAS:GCaMP6s); Tg(sst2:RFP); Tg(ins:mCardinal)* donor embryos were injected with 200 pg *sox32* mRNA, generated with mMESSAGE mMACHINE ™ SP6 transcription kit (Thermo), at the one-cell-stage to enhance endoderm formation. *Tg(elavl3:GAL4-VP16); Tg(UAS:BoTxBLC-GFP)* host embryos were injected with 0.3 ng *sox32* morpholino (Gene Tools LLC) at the one-cell-stage to deplete endoderm formation. At the 1000-cell-stage, single cells were removed from donor embryos with a glass capillary needle, and ~40 cells were transplanted into host embryos in the region fated to become endoderm-derived tissues. Only larvae that displayed complete endoderm transplant, without obvious defects, and carried the relevant transgenes were used for downstream imaging experiments. Control chimeras were negative for BoTxBLC-GFP expression.

### In vivo confocal microscopy

Live zebrafish between 100-110 hours post fertilization were anesthetized with 0.015% Tricaine and mounted in 0.8% low melting agarose in egg water containing 0.005% Tricaine for confocal imaging. Zeiss LSM880 upright laser scanning confocal microscopes equipped with a Plan-Apochromat 20x/NA1.0 dipping lens was used for imaging. Time-lapse calcium imaging was conducted in 25°C conditions and z-stacks were taken at 5 s intervals for 1-2 hr. Larvae were exposed to 75 mM glucose containing egg water at the indicated time points during the time-lapse imaging to increase whole larvae glucose levels within a physiological range (Figure 2 – Fig. Suppl. 1). For photo-inhibition experiments, GCaMP6s signal was visualized with a Chameleon Vision II Ti:Sapphire Laser (Coherent) laser at 920 nm. For all other experiments, GCaMP6s signal was visualized with an Argon laser at 488 nm.

### Two-photon laser ablation

A Chameleon Vision II Ti:Sapphire Laser (Coherent) mounted on a Zeiss LSM880 microscope was used for two-photon single-cell laser ablation of peri-islet neurons labelled by the *Et(1121A:GAL4FF)* enhancer trap. The tunable laser was set at 800 nm to scan an ablation area of 4 μm^2^ at a scan speed of 1 with 10 iterations. Following ablation, embryos were rested for 5 min before continuation of calcium imaging. Controls were mock ablation of cells within 20 μm from the peri-islet neurons. Animals from the same clutch were randomly assigned to ablation and control groups.

### Photo-inhibition of nerve activity

All experiments with *Tg(elavl3:sypb-miniSOG2-P2A-mScarlet)* were done on F1 to F3 larvae displaying strong and uniform mScarlet signal. Controls were transgene-negative siblings. For assessing swimming behavior, 110 hpf larvae were individually isolated in 9 mm diameter circular wells filled with 200 μl egg water. A Nikon SMZ25 stereomicroscope with a P2-SHR PlanApo 1x/NA0.15 objective was used for time-lapse imaging of larvae at 1 s intervals, the same animals were imaged pre-light and post exposure to a 5 min pulse of blue light (3.8 mW LED, 466/40 filter, Lumencor Sola Light Engine). For assessing changes in glucose levels, pools of 40 larvae swimming in 9 cm diameter petri-dishes filled with 35 mL egg water were exposed to 0.3 mW blue LED light for 30 min prior to sample collection, control animals were not exposed to blue light and were randomly selected siblings from the same clutch. Notably, acute blue light exposure alone mildly reduced swimming behaviour and increased glucose levels (Figure 4 – Fig. Suppl. 1C, Figure 4A). Induction of a stress response likely led to this increase in glucose levels in wild-type larvae, as chronic 14 hr exposure to blue LED light for 3 consecutive days kills zebrafish larvae^53^. For assessing calcium changes, larvae were exposed to a 3 min pulse of blue light (2.1 mW 470 nm LED, Colibri) and GCaMP6s signal was measured pre- and post-blue light exposure. While blinding was not possible, we randomized the order in which animals on a given day were imaged.

### Wholemount immunostaining

Zebrafish were euthanized with tricaine overdose prior to overnight fixation in 4% paraformaldehyde dissolved in PBS containing 120 μM CaCl_2_ and 4% sucrose, pH7.4. The skin was manually removed with forceps, without disturbing the internal organs and the zebrafish were permeabilized with 1% TritonX-100, 1% DMSO containing PBS for 3 h at room temperature for most staining protocols. For cx43 staining, antigen retrieval with 150 mM Tris-HCl pH 9 at 70°C for 15 min and acetone permeabilization at −20°C for 20 min was performed. Following blocking with 5% donkey serum (Jackson Immunoresearch) in blocking buffer (Dako), samples were incubated in primary antibodies overnight at 4°C, washed 4x with 0.025% TritonX-100 containing PBS, incubated in secondary antibodies overnight at 4°C, washed 4x, incubated in an increasing glycerol gradient of 25%, 50%, and 75%, and mounted in VectorShield mounting medium. The following antibodies and dilutions were used: mouse anti-glucagon (1:200, Sigma G2654), chicken anti-GFP (1:200, Aves GFP-1020), mouse anti-acetylated Tubulin (1:200, Sigma T7451), rabbit anti-connexin 43 (1:100, Sigma C6219). Secondary antibodies used in this study include donkey anti-rabbit AlexaFluor568 (1:400, Thermo A10042) and AlexaFluor488 (1:300, Jackson 711-545-152), donkey anti-mouse AlexaFluor488 (1:300, Jackson 715-545-150) and AlexaFluor647 (1:200, Jackson 715-605-150), donkey anti-chicken AlexaFluor488 (1:300, Jackson 703-545-155). Nuclei were stained with 25 μg/ml DAPI (Sigma). Images were taken on Zeiss LSM880 or LSM800 laser scanning confocal microscopes equipped with a 25x/NA0.8 objective.

### Data analysis

Image data were analyzed using Imaris (Bitplane) and Fiji (ImageJ). Correlation and fraction time analysis was performed using Matlab (code available upon request). Statistical analysis was performed using Prism 8 (GraphPad). D’Agostino & Pearson test was used to assess Gaussian distribution of the data and subsequent selection of parametric or nonparametric tests. For comparison between two groups, a two-tailed Student’s *t*-test or a Mann-Whitney test was used, respectively, to determine the p-values. For comparison between more than two groups, an ordinary one-way ANOVA with Holm-Sidak’s multiple comparisons test or a Kruskal-Wallis test with Dunn’s multiple comparisons test was used, respectively, to determine the p-values. The number of animals or cells analyzed, and the p-values are reported in the figure and figure legends. All experiments were repeated on different days using at least three different clutches of animals.

## Acknowledgments

These studies were supported by funds from the Max Planck Society to D.Y.R.S. Y.H.C.Y was supported by CIHR Postdoctoral Fellowships, an HFSP Long-Term Fellowship, an EMBO Long-Term Fellowship, and NIG-JOINT funding. L.J.B.B. was supported by a Sir Henry Wellcome Postdoctoral Fellowship. K.K. was supported by an NBRP grant from AMED. We thank Arica Beisaw for the plasmid with *P2A-mScarlet* and James Johnson for critical reading of the manuscript. The referenced *cx43* mutant line was generously provided by Matthew Harris.

## Author contributions

Y.H.C.Y. conceived the study, designed/performed the research, analyzed data, and wrote the manuscript. L.J.B.B. analyzed data, provided critical feedback and edited the manuscript. C.R., S.T.J. and H.M.M. designed/performed the research. K.K. provided the *Et(1121A:GAL4FF)* and *Tg(UAS:GCaMP6s)* lines and edited the manuscript. D.Y.R.S. provided the resources and critical feedback, and edited the manuscript.

---

**Figure 2 - Figure Supplement 1.**
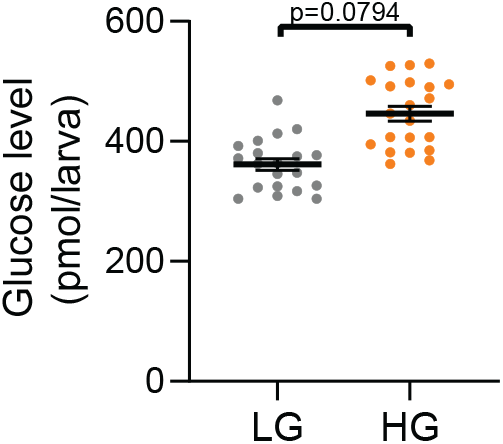
Glucose treatment elevates whole larva glucose levels. Zebrafish at 100 hpf were treated with 75 mM glucose in egg water for 1 hr. Whole larva free glucose level measurements at 101 hpf, mean ± SEM, n=11-19 batches of 5 larvae per replicate.

**Figure 4 - Figure Supplement 1.**
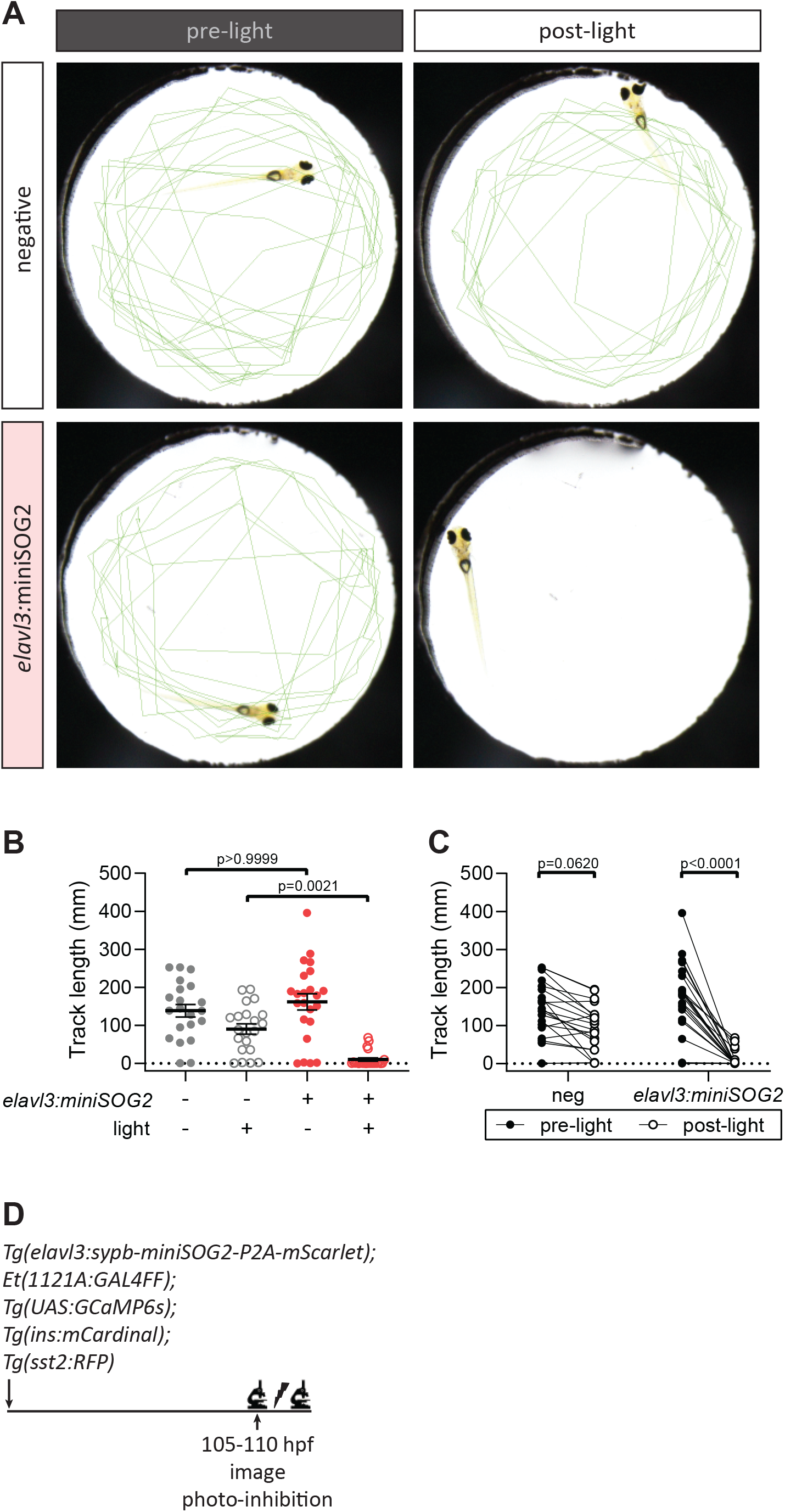
Swimming activity is reduced following blue light exposure in *Tg(elavl3:sypb-miniSOG2-P2A-mScarlet)* larvae. **A.** Swimming activity of *Tg(elavl3:sypb-miniSOG2-P2A-mScarlet)* zebrafish at 110 hpf was tracked pre- and post-blue light exposure for 5 min. Green lines show tracks at the end of the time-lapse. **B-C.** Quantification of swimming activity of *Tg(elavl3:sypb-miniSOG2-P2A-mScarlet)* zebrafish at 110 hpf, mean ± S.E.M., n=21-23 animals, p-values from ANOVA and t-tests, respectively. **D.** Schematic of optogenetic neural inhibition experiment.

**Figure 5 - Figure Supplement 1.**
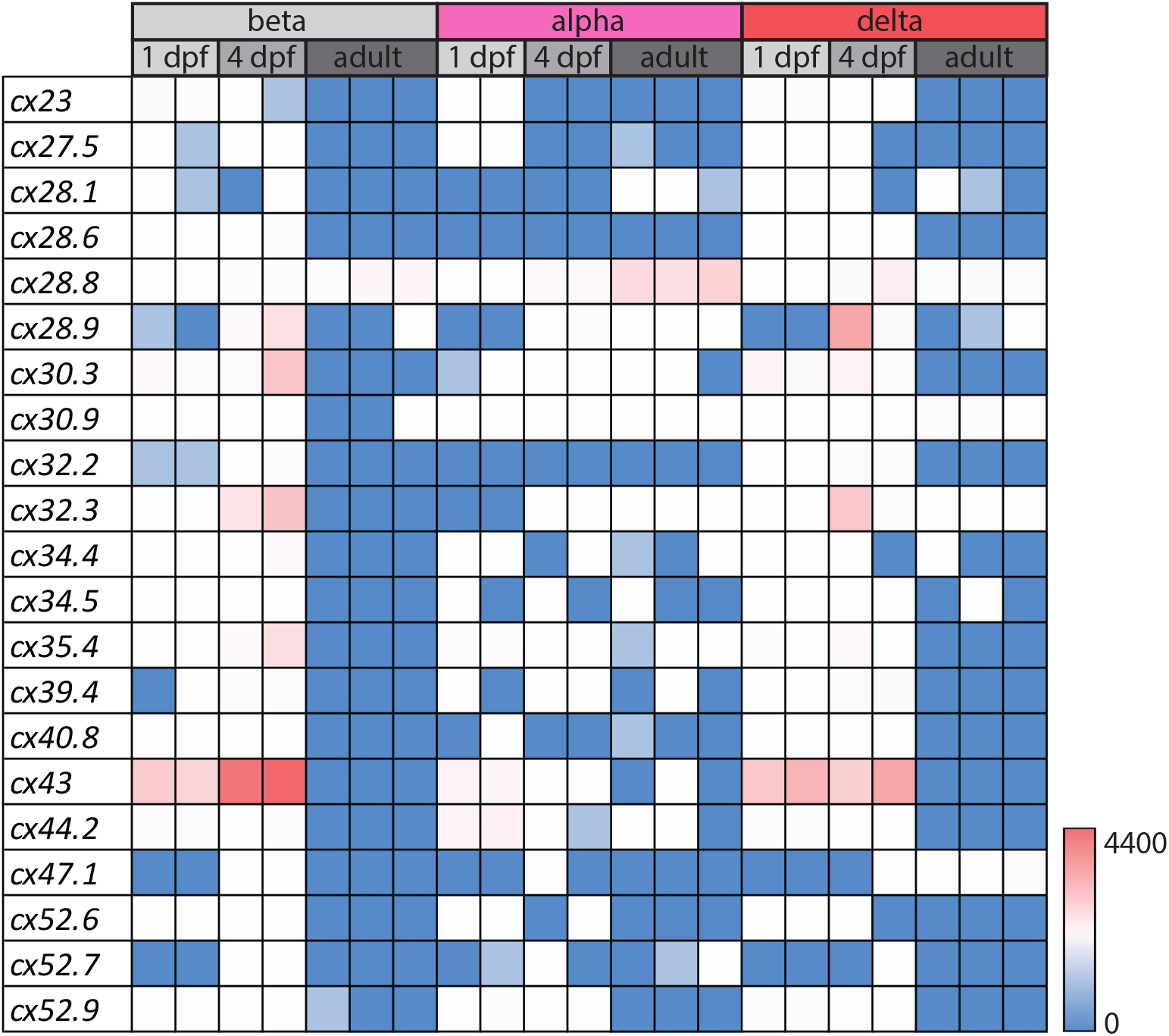
Expression of gap junction genes in FACS sorted zebrafish islet cells. Expression profile of gap junction genes from in-house and published RNAseq datasets of FACS sorted zebrafish pancreatic beta, alpha, and delta cells at embryonic (24-28 hpf), early larval (96-100 hpf), and adult stages^37^.

**Figure 5 - Figure Supplement 2.**
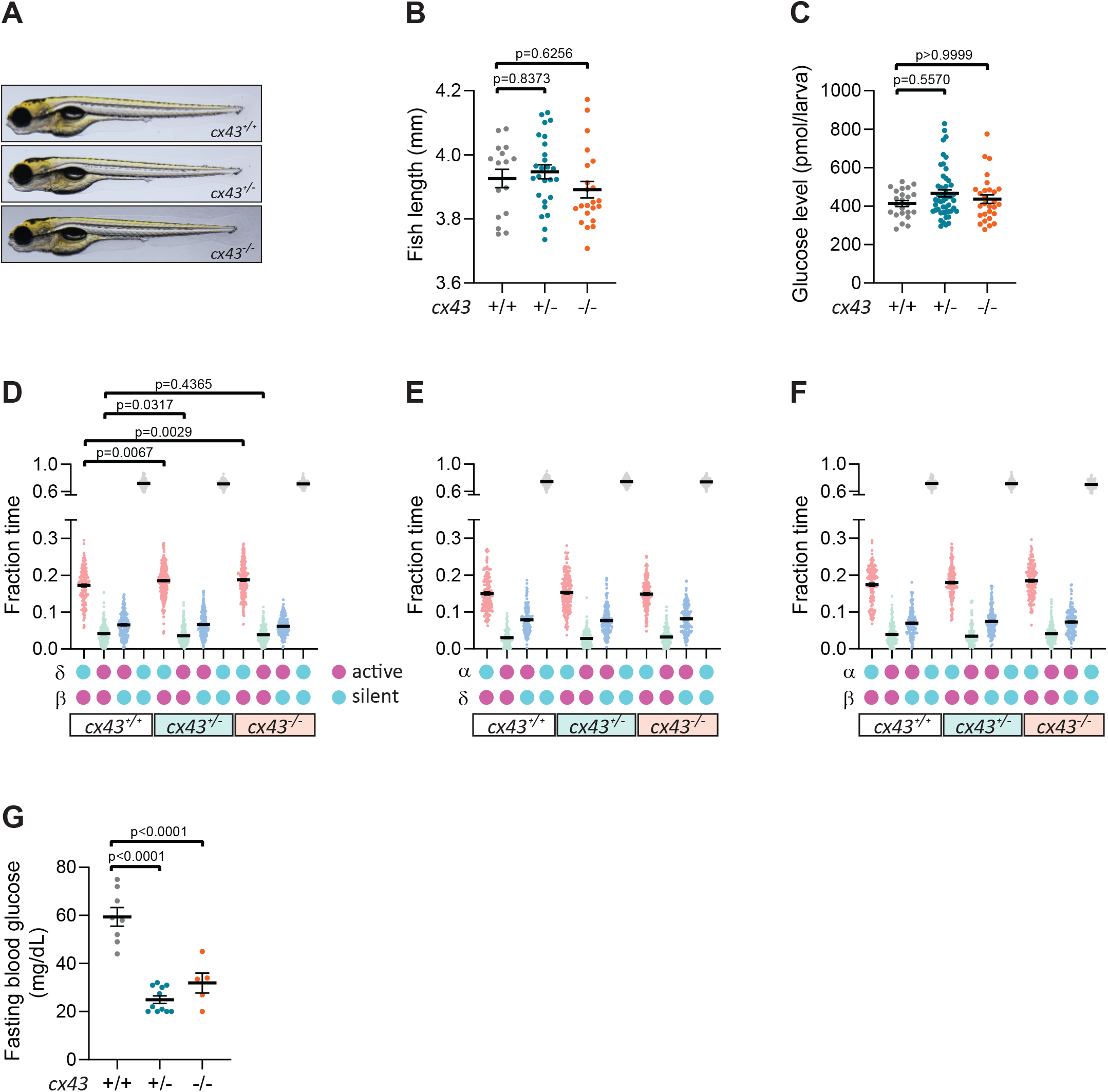
Changes in heterotypic interactions between pancreatic islet cells in *cx43* mutants. **A.** *cx43^+/+^, cx43^+/−^*, and *cx43^−;/−^* larvae imaged at 100 hpf. **B.** Body length measurements at 100 hpf, mean ± S.E.M., n=18-26 animals. **C.** Whole larva free glucose level measurements at 100 hpf, mean ± S.E.M., n=23-51 batches of 5 larvae per replicate. **D-F.** Fraction time analysis of heterotypic delta-beta **(D)**, alpha-delta **(E)** and alpha-beta **(F)** cell pairs, mean ± SEM, n=165-293 cell pairs; magenta circle, active state; cyan circle, silent state. **G.** Fasting blood glucose levels in adults, mean ± SEM, n=5-11 animals, p-values from ANOVA.

**Figure 1 – Movie Supplement 1.** 100 hpf *Et(1121A:GAL4FF); Tg(UAS:GCaMP6s); Tg(ins:mCardinal); Tg(sst2:RFP)* zebrafish stained with DAPI (DNA).

**Figure 1 – Movie Supplement 2.** Intracellular calcium dynamics in pancreatic islet cells (including delta, beta, alpha, and unidentified cells) in 100 hpf *Et(1121A:GAL4FF); Tg(UAS:GCaMP6s); Tg(ins:mCardinal); Tg(sst2:RFP)* zebrafish.

**Figure 5 – Movie Supplement 1.** Intracellular calcium dynamics in pancreatic islet cells in *Et(1121A:GAL4FF); Tg(UAS:GCaMP6s); Tg(ins:mCardinal); Tg(sst2:RFP) cx43^+/+^*, *cx43^+/−^,* and *cx43^−;/−^* larvae imaged at 100 hpf. Normalized calcium traces of beta cells mapped to their location within the pancreatic islet.

## Graphical Abstract

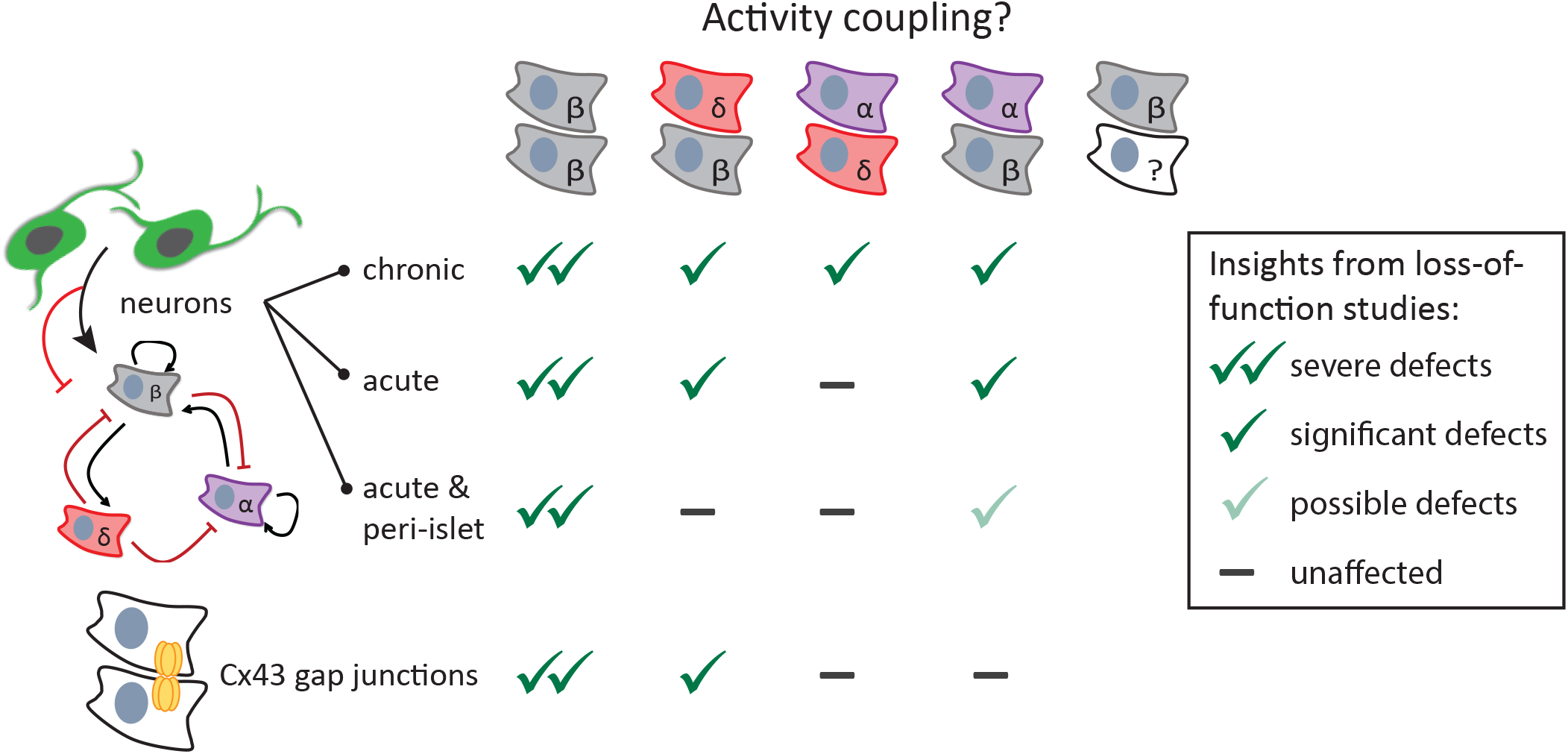

